# Temporal Information Encoding in Isolated Cortical Networks

**DOI:** 10.1101/2024.09.25.614992

**Authors:** Zubayer Ibne Ferdous, Yevgeny Berdichevsky

## Abstract

Time-dependent features are present in many sensory stimuli. In the sensory cortices, timing features of stimuli are represented by spatial as well as temporal code. A potential mechanism by which cortical neuronal networks perform temporal-to-spatial conversion is ‘reservoir computing’. The state of a recurrently-connected network (reservoir) represents not only the current stimulus, or input, but also prior inputs. In this experimental study, we determined whether the state of an isolated cortical network could be used to accurately determine the timing of occurrence of an input pattern – or, in other words, to convert temporal input features into spatial state of the network. We used an experimental system based on patterned optogenetic stimulation of dissociated primary rat cortical cultures, and read out activity via fluorescent calcium indicator. We delivered input sequences of patterns such that a pattern of interest occurred at different times. We developed a readout function for network state based on a support vector machine (SVM) with recursive feature elimination and custom error correcting output code. We found that the state of these experimental networks contained information about inputs for at least 900 msec. Timing of input pattern occurrence was determined with 100 msec precision. Accurate classification required many neurons, suggesting that timing information was encoded via population code. Trajectory of network state was largely determined by spatial features of the stimulus, with temporal features having a more subtle effect. Local reservoir computation may be a plausible mechanism for temporal/spatial code conversion that occurs in sensory cortices.

**Significance Statement:** Handling of temporal and spatial stimulus features is fundamental to the ability of sensory cortices to process information. Reservoir computation has been proposed as a mechanism for temporal-to-spatial conversion that occurs in the sensory cortices. Furthermore, reservoirs of biological, living neurons have been proposed as building blocks for machine learning applications such as speech recognition and other time-series processing. In this work, we demonstrated that living neuron reservoirs, composed of recurrently connected cortical neurons, can carry out temporal-spatial conversion with sufficient accuracy and at sufficiently long time scale to be a plausible model for information processing in sensory cortices, and to have potential computational applications.

## Introduction

Time-dependent features are present in many sensory stimuli, including moving visual objects, speech, or a fingertip running over a textured surface. Temporal code, based on ‘when’ a neuron activates, is initially used by the nervous system to represent time-dependent features. In the sensory cortices, time features are also represented by spatial code, based on ‘which’ neuron is activated (1–3). A cortical neuron may be preferentially activated by a specific sequence of stimuli. Spatial code formed by the pattern of activation of these neurons represents information that was originally temporally encoded (4). One proposed mechanism of temporal-to-spatial conversion is the formation of a ‘computational reservoir’ (sometimes called liquid-state machine) by neurons (5). A network of recurrently connected neurons represents a system (reservoir) with a state that depends on previous state of the reservoir as well as the current inputs (6). State of the reservoir is defined by activation of neurons within the reservoir, and can be visible (activation corresponds to neuronal firing rates) or hidden (activation of neuronal dynamics such as long-lasting currents or short-term synaptic plasticity) (7). In reservoir computing paradigm, time-varying *u*(*t*), representing inputs *u*_1_(*t*), *u*_2_(*t*), …., *u*_N_(*t*), is continuously delivered to the reservoir *r*. Reservoir state at time t is *x*(*t*), representing states of individual reservoir elements *x*_1_(*t*), *x*_2_(*t*) …. *x*_M_(*t*). Reservoir is recurrently connected such that reservoir state at time *t* + 1 is: *x*(*t*+1) = *r*(*u*(*t*), *x*(*t*)), where *r* is the function carried out by the reservoir elements. Due to its recurrent connections, the reservoir is capable of retaining information about input *u* at preceding values of time by encoding it in its state *x*. Recurrent connections also enable communication between different reservoir elements. Information may also be retained in the neuronal dynamics. Thus, state *x*(t) depends both on temporal information in the input (*u* has different values at times … *t*-2, *t*-1, and *t*) and spatial information in the input (*u*_1_(*t*), *u*_2_(*t*), … *u*_N_(*t*) have different values). Memoryless readout map f can be used to transform the reservoir state *x* into an output *y*: *y* = f(*x*(*t*)). Such readout map may represent a higher order cortex that makes use of the spatiotemporally-integrated information encoded in reservoir state *x*(*t*).

The ability of reservoir computing to integrate temporal as well as spatial information has led to its use in machine learning applications such as recognition of speech (8, 9) and continuous gestures (10). High performance of reservoirs composed of artificial neurons in turn motivated investigation into reservoirs composed of biological, living neurons. Investigations focused on two related objectives: gaining a better understanding of the role that a reservoir-like recurrently connected network of neurons may play in spatiotemporal information processing in the brain, and harnessing the ability of recurrently connected networks of neurons to perform computational tasks. Use of biological neurons to form computational reservoirs may increase energy efficiency of computation – a metric that is becoming increasingly important in the design of computing systems (11). Experimental reservoirs have been constructed from dissociated rat cortical neurons, as well as neurons differentiated from human induced pluripotent stem cells. These systems demonstrated the ability of reservoirs composed of living neurons to differentiate between spatial stimulus patterns with high accuracy (12). Differentiation of spatiotemporal sequences, based on analysis of neuronal activity at one point in time (i.e. using purely spatial coding to classify inputs with temporal as well as spatial features) was also successful, albeit at a relatively low accuracy (13). Experimental neuronal reservoirs were used for speech recognition by converting spoken digits into spatiotemporal input patterns, and classifying the reservoir outputs (14). Human organoid-based reservoirs were used in a similar fashion to classify different speakers (15). These studies demonstrated that reservoirs composed of recurrently connected neurons were capable of classifying input patterns that contained mixed temporal and spatial encoding. However, the ability of these reservoirs to convert temporal input features into spatial code has not yet been shown. It is also not clear whether reservoirs can carry out this conversion with accuracy that is sufficient for processing of sensory input information, or that would be practical for computational systems. In this work, we aimed to answer these questions. We experimentally show that spatial pattern of neuronal activation in a reservoir (reservoir state *x*(t)) embeds temporal features of the input (*u* at times from *t*-T to *t*) that can then be accurately extracted by function f.

## Results

### Optical Interface for Neuronal Reservoirs

We devised an all-optical setup to stimulate neurons in confined cortical cultures and detect evoked activity at the same time. Dissociated cortical neurons were placed into a well cut in a polydimethylsiloxane (PDMS) film to ensure that a significant proportion of the network was optically accessible (Fig. 1A(i)). We co-infected these neurons with channelrhodopsin 2 (ChR2) and jRGECO1a (red fluorescent genetically encoded calcium indicator) (16). By day 14 in vitro (DIV 14), confined neurons formed a synaptically-connected network, and most neurons expressed ChR2 and jRGECO1a (Fig. 1B). We then placed dishes containing these confined networks in a mini-incubator on a stage of an inverted dual-light path microscope. We used a digital micromirror device-based pattern stimulator to deliver pulses of blue light to predefined regions of the network (Fig. 1A(ii)). Minimum stimulation parameters (light spot size, power, and pulse duration) that reliably evoked action potentials in illuminated neurons were determined (Fig. S1). Light patterns in subsequent experiments were designed to match or exceeded these minimal parameter settings. We then optimized excitation light delivery for observing changes in jRGECO1a fluorescence. Peak excitation wavelength for jRGECO1a was reported as 550 nm (16). We observed 11± 5 mV depolarization when 550 nm excitation was delivered to neurons expressing both jRGECO1a and ChR2, due to partial excitation of ChR2 at this wavelength (Fig. S2A). We red-shifted the excitation to 580 nm to eliminate this undesirable depolarization at 5% intensity level, which was used for subsequent experiments (Fig. S2B). At these settings, an action potential in a neuron resulted in approximately 1% change in jRGECO1a fluorescence over baseline (Δ*F*/*F*) (Fig. S3). Parallel delivery of ChR2 excitation patterns and wide-field jRGECO1a excitation light enabled us to simultaneously stimulate and record neuronal activity in cultured cortical neuron networks.

**Figure 1.**
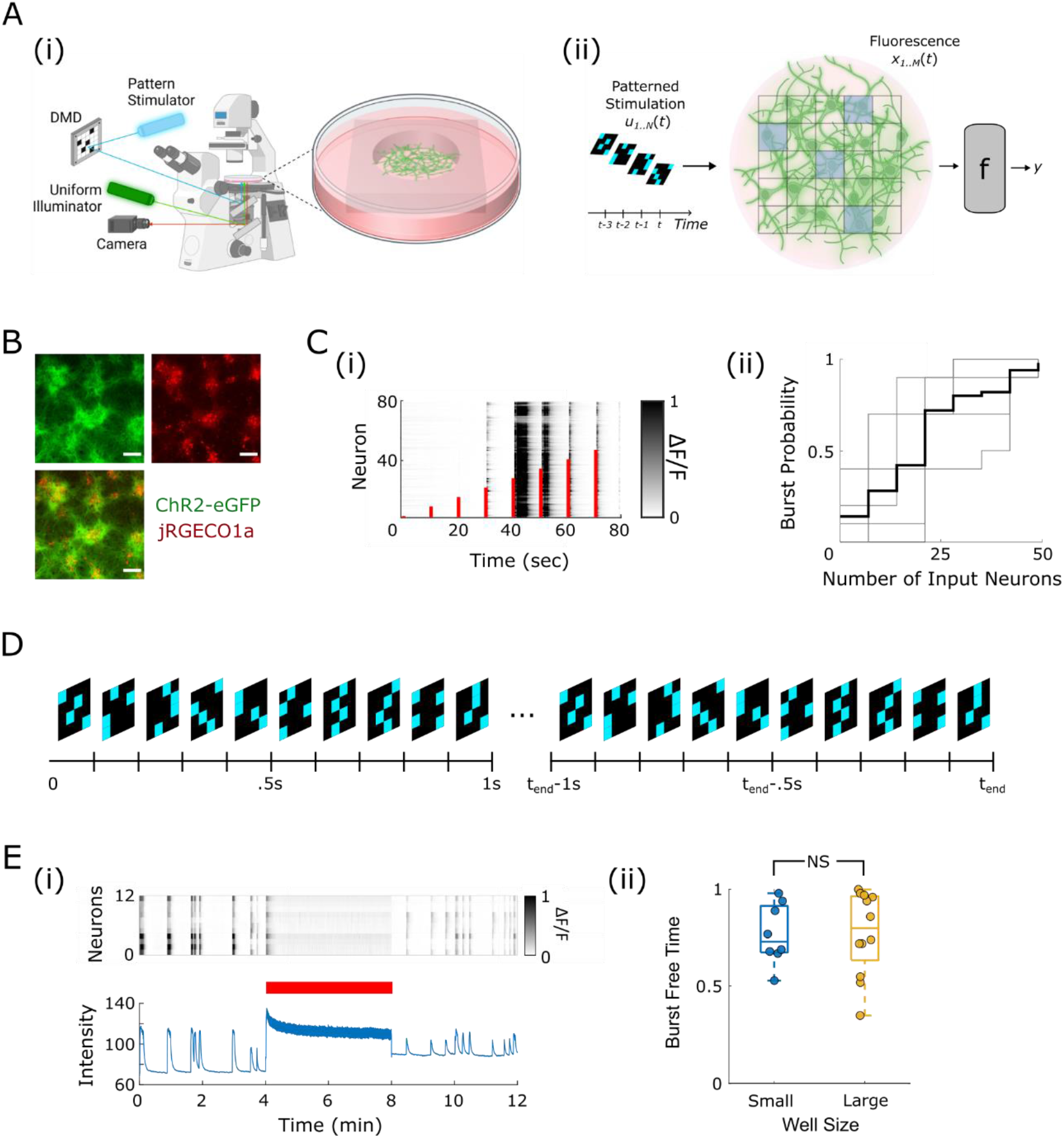
Optical stimulation and detection of neuronal activity. (**A**) (i) Schematic of all-optical experimental platform. (ii) Schematic of the reservoir computing paradigm using neuronal cultures with an optical interface. *N* = number of input grids, *M* = number of output neurons. (**B**) Fluorescence images of jRGECO1a and ChR2 expression. (**C**) (i) Plot of ΔF/F signals from multiple neurons while network was subjected to episodic stimulation. Stimuli times are indicated by red lines, length of the red lines corresponds to the number of optically stimulated (input) neurons. (ii) Plot of burst probability per number of input neurons for *n* = 5 cultures. Data for individual cultures is gray, average is shown as a thick black line. (**D**) Continuous input pattern delivery; illuminated grid cells are shown in blue. Repeating sequence of 10 patterns is 1 sec long. (**E**) Burst-free stimulation: (i) Representative raster plot and average jRGECO1a fluorescence intensity before, during, and after continuous input pattern delivery (indicated with red line); (ii) Burst-free time in cultures in small (*n* = 8) and large (*n* = 12) wells was not significantly different (*p* = 0.94, two-sample *t*-test).

### Continuous Stimulation

Cultured networks of dissociated cortical networks fire spontaneous population bursts of different duration (17). These bursts are a type of spontaneous activity involving synchrony between large numbers of neurons, and reminiscent of early developmental activity (18). Spontaneous population bursts interfere with information processing in cultured networks (13, 19). Episodic stimulation of cultured networks can also evoke undesirable population bursts (12). We quantified evoked bursts in our cortical cultures, containing an average of 567 neurons per culture (*n* = 5 cultures). We activated 1-50 input neurons with a step of 7 inputs (Fig. 2C(i)). Stimulation was delivered at 0.1 Hz using the minimum reliable stimulation parameters and each input pattern was repeated 10 times. Population bursts were defined as events where at least 80% of output neurons had ΔF/F > 0.2. Reliability of population burst evocation increased with the number of stimulated (input) neurons and reached 100% for all 5 cultures with 50 input neurons (i.e. stimulation of <10 % of neurons in the network) (Fig. 2C(ii)). This type of evoked response – activation of nearly all neurons in the network due to a sparse input – is substantially different from the sparse neuronal activation of an intact cortex by sensory input. We therefore decided to develop a stimulation protocol that resulted in a sparser activation without population bursts. In earlier work, Wagenaar and colleagues (20) showed that distributed and continuous, as opposed to episodic, stimulation of cultured networks suppressed population bursts. This was likely due to activation of internal neuronal dynamics such as afterhyperpolarization currents and short-term synaptic depression. These dynamics may be active in an awake cortex, but inactive in cultured neurons with no sensory input. We reasoned that if we designed continuous stimulation such that a significant number of neurons in the cultured network were activated frequently, neurons in the cultured network may have more awake-cortex like dynamics, and be less susceptible to synchronized bursting. We developed a protocol where a sequence of patterns is continuously delivered to the cultured network. Patterns were designed by dividing the field of view into a rectangular grid, and delivering light pulses to a subset of the grid cells for each pattern (Fig. 1D). Grid spacing was varied. Illumination for each pattern consisted of a 50 Hz train of five pulses of 10 msec pulse duration (these parameters were used to achieve maximum evoked response with minimum train duration, Fig. S4). Delivery of each pattern took 100 msec, with a full sequence of 10 patterns delivered every second. Neurons were designated as ‘input’ neurons if directly optically stimulated, and as ‘output’ neurons if never directly stimulated. Pattern sequences were designed such that each input neuron was stimulated at least once per second. Proportion of input neurons to total number of neurons in the network was > 30%; this was possible due to limiting the physical extent of the network via confinement in PDMS well. Continuous delivery of pattern sequences resulted in suppression of spontaneous and evoked population bursts (Fig. 1E). There were no significant differences between burst-free time in cultures confined in small (1 mm^2^) and large (4 mm^2^) PDMS wells. We used large wells in subsequent experiments as they contained more output neurons for use in classification. Stimulation protocol enabled both spatial and temporal input encoding: spatial information was contained in the pattern design, whereas temporal information was contained in pattern sequence.

**Figure 2.**
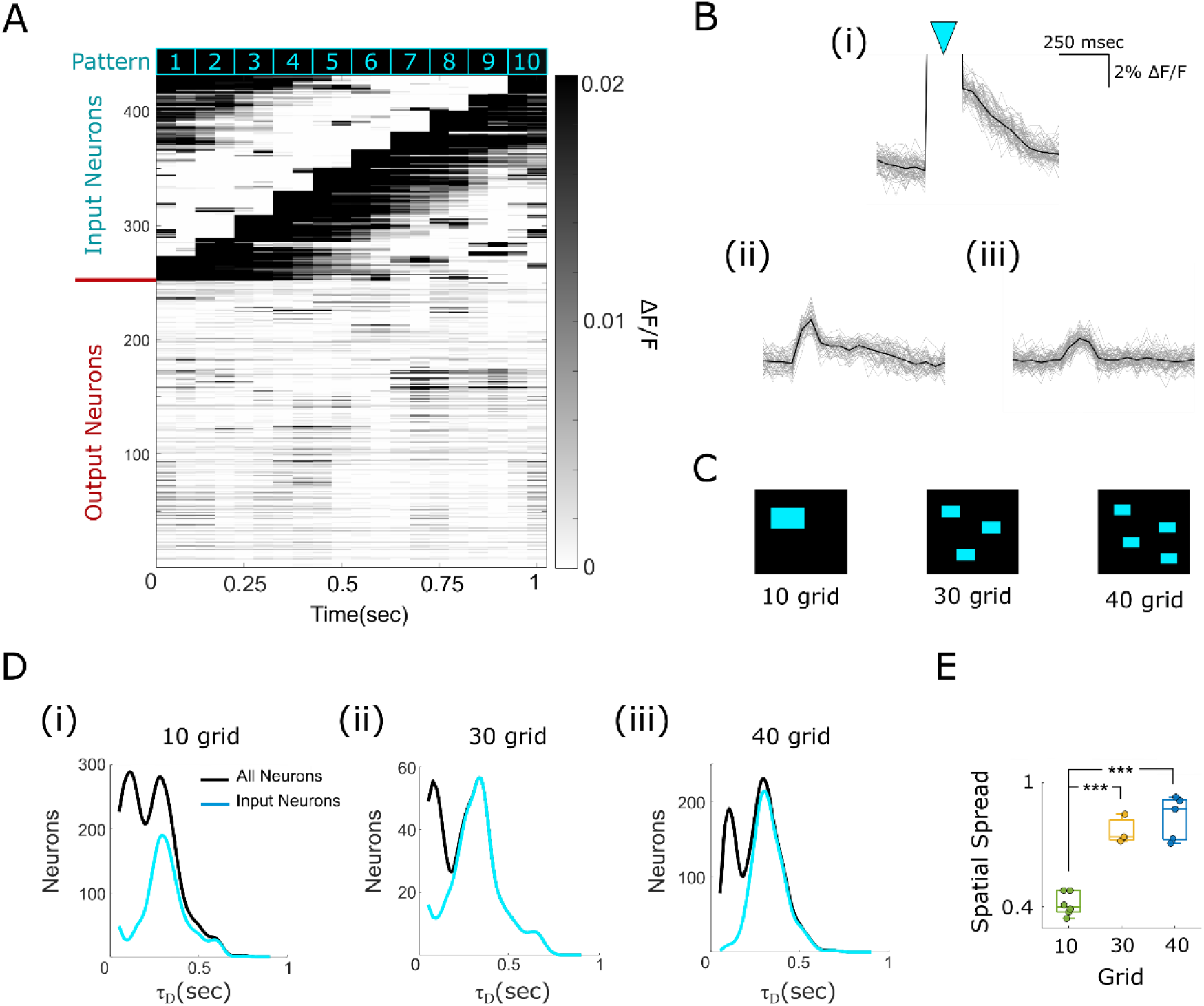
Activity evoked by different stimulation designs. (**A**) Representative raster plot of average activity evoked by a sequence of 10 patterns, (**B**) Representative traces of activity evoked in (i) input neurons (blue arrowhead indicates stimulus artifact), and (ii, iii) output neurons. Responses to a single stimulus sequence are shown as gray traces, and average response to n = 50 sequence repeats is shown as a black trace. Output neuron in (ii) has a longer time decay constant (*τ*_D_) than output neuron in (iii). (**C**) Different grid designs. (**D**) (i-iii) Histograms of *τ*_D_ in cultures stimulated with patterns based on different sequences. *n* = 2036 neurons in 6 cultures, *n* = 407 neurons in 3 cultures, and *n* = 1408 neurons in 5 cultures for ‘10 grid’, ‘30 grid’, and ‘40 grid’, respectively. (**E**) Spatial spread in cultures with ‘10 grid’ (*n* = 6 cultures), ‘30 grid’ (*n* = 3 cultures), and ‘40 grid’ (*n* = 5 cultures) stimulus patterns. *** p < 0.001, two-sample *t*-test.

### Evoked Activity

Stimulation patterns were designed by dividing the field of view into grids of different spacing, and delivering light to 1, 3, or 4 grid cells per pattern. These pattern designs were termed ‘10 grid’ (1 grid cell per pattern, largest cell size), ‘30 grid’ (3 grid cells per pattern, medium cell size), and ‘40 grid’ (4 grid cells per pattern, largest cell size) (Fig. 2(C)). Patterns were designed so that approximately 40% of neurons functioned as ‘input neurons’, regardless of the total number of neurons (Fig. S5). Sequences of 10 different patterns were continuously delivered between 40 and 100 times. Representative examples of evoked activity in input and output neurons are shown in Fig. 2A, B. We noticed that evoked activity in input neurons was characterized by the presence of a stimulus artifact (Fig. 2 B(i)), due to leakage of stimulation light into detection filter, and to activation of jRGECO1a by blue light. We excluded the stimulus artifact in subsequent analysis. After stimulus, input and some output neurons (Fig. 2B(i, ii)) were characterized by long decay time constants (τ_D_) of evoked fluorescence changes, but most of evoked activity in output neurons had short decay constants (Fig. 2B(iii)). We noticed that spatial spread and decay of evoked activity strongly depended on the grid design. Evoked activity decayed faster in output neurons of networks stimulated with ‘30 grid’ and ‘40 grid’ patterns compared to ‘10 grid’ patterns (Fig. 2D). Spatial spread of evoked activity in output neurons was significantly larger due to ‘30 grid’ and ‘40 grid’ stimulation compared to ‘10 grid’ stimulation (Fig. 2E). We selected ‘40 grid’ stimulation for subsequent experiments since evoked activity was spread through a larger portion of the network than for ‘10 grid’ stimulation, and it offered more design freedom for patterns than ‘30 grid’ stimulation.

### Pattern Selectivity

Examination of evoked activity during stimulation with a sequence of patterns revealed preferential activation of individual neurons during specific patterns (pattern selectivity) (Fig. 3A). Approximately 49% of output neurons (533 out of 1090) in 5 cultures exhibited significant pattern selectivity (Fig. 3B). We then examined whether activity of output neuron population was separable in low dimensional space by each pattern in the stimulus sequence. We used Principal Component Analysis (PCA) on population vectors of output neuron activity, using only neurons with significant pattern selectivity. Activity during last 50 msec of a given pattern during stimulation sequence was used to construct a population vector for that pattern. Scores on first 3 principal components (PCs) clustered by stimulation pattern (Fig. 3C(i)). We calculated silhouette values for each point in pattern-based clusters to determine degree of clustering. Silhouette values higher than zero indicate good cluster matching, whereas values lower than zero indicate poor matching. For culture shown in Fig. 3C(i), most evoked responses were well clustered by pattern. We calculated average silhouette values for each of 10 patterns in 5 cultures, and found that they were significantly higher than zero (Fig. 3C(iii)). This demonstrated that response of output neuron population to a sequence of patterns was pattern-selective.

**Figure 3.**
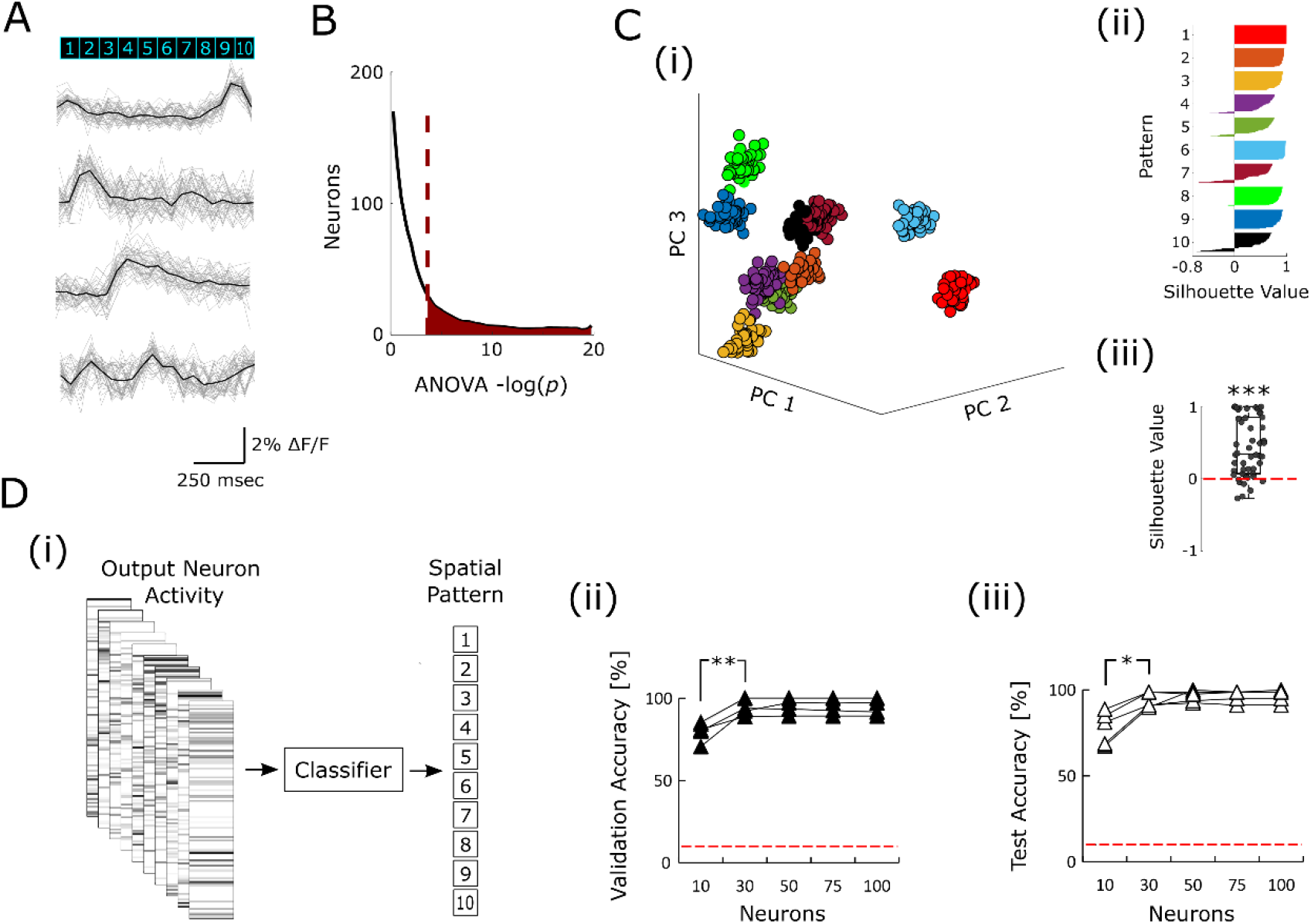
Pattern selectivity. (**A**) Representative traces of evoked activity of 4 output neurons during stimulation with a sequence of 10 patterns. Activity evoked by *n* = 40 single sequences (gray) and average activity (black) are shown. (**B**) Histogram of *p* values obtained with one-way ANOVA on evoked responses of *n* = 1090 output neurons (5 cultures) to 10 different patterns in a repeating sequence. Horizontal axis is logarithmic. Significance threshold was determined by dividing *p* = 0.05 by average number of output neurons per culture (*p* = 2.3*10^−4^), and indicated by a red dashed line on the graph. Neurons with -log(*p*) > 20 are not shown. (**C**) (i), PCA scores of evoked activity in output neuron population on first 3 PCs. Each dot represents response to a single pattern stimulation. Dots are color coded by pattern. (ii), Silhouette values of each response, relative to clusters assigned by pattern. Color coding is same in (i) and (ii). (iii), Silhouette values for each pattern in each of 5 cultures (*n* = 50). *** *p* < 0.001 on two-sided Wilcoxon signed rank test. (**D**) (i) Spatial pattern inference: population vectors of output neuron activity are classified to 10 stimulation patterns. (ii) validation and (iii) testing classification accuracy achieved using varying numbers of output neurons to construct population vectors. Random classification chance is 1/10 = 10%, indicated by red dashed line. Data is for *n* = 5 cultures. * *p* = 0.011, ** *p* = 0.003 on two-sample *t*-test.

*Pattern Inference*

We next checked whether evoked activity of output neuron population could be used to accurately infer the stimulation pattern. Population activity vectors were constructed as before, but using only neurons with lowest *p*-values on one-way ANOVA for pattern selectivity. Observations (population vectors from 40 sequence repeats) were divided into training (80%) and testing (20%) datasets. Support Vector Machine (SVM) classification model was trained and then tested using these data sets (Fig. 3D). SVM model was able to classify population vectors by pattern with 95% average validation accuracy (using training dataset, Fig. 3D(ii)) and 94% average testing accuracy (using testing dataset, Fig. 3D(iii). These accuracies were achieved using only 30 output neurons; use of higher number of output neurons did not lead to an increase in classification accuracy. We conclude that high specificity of output neurons to spatial stimulation patterns results in relatively few output neurons (compared to total) required to infer stimulation pattern with high level of accuracy.

### Temporal Sequence Inference

We then examined whether activity of output neurons contained information about temporal history of stimuli. We designed an experiment where 5 different temporal sequences of 10 spatial stimulus patterns each were delivered to the input neurons. Different sequences contained the same 10 patterns, but in different order (Fig. 4A). Output activity was examined during patterns 6 and 10. Pattern order was designed such that pattern 1 was delivered at times ranging from 100 msec to 900 msec relative to pattern 10 (Fig. 4A(i)), and from 500 msec to 800 msec relative to pattern 6 (Fig. 4A(ii)). The remainder of pattern order was identical between sequences. This design enabled us to determine whether activity of output neurons reflected the temporal position pattern 1, as late as 900 msec and 9 stimulus patterns after pattern 1 was delivered. The 5 sequences were delivered repeatedly (40 – 50 trials in each of 5 cultures, one trial is shown in Fig. 3B). We analyzed activity of 1090 output neurons in 5 cultures during stimulus patterns 6 and 10, and found that only 9 of the 2180 analyzed activities (0.4 %) were significantly different between sequences (Fig. 4C). We then constructed output neuron population activity vectors during patterns 6 and 10, per sequence, and reduced their dimensions to 3 principal components using PCA. Vector scores on these components did not visually cluster by sequence (Fig. 4D(i)). Silhouette values for putative sequence-based clusters were significantly below zero (Fig. 4D(ii,iii)). This indicated poor or non-existent clustering by temporal sequence, in contrast to good clustering by spatial pattern (Fig. 3C).

**Figure 4.**
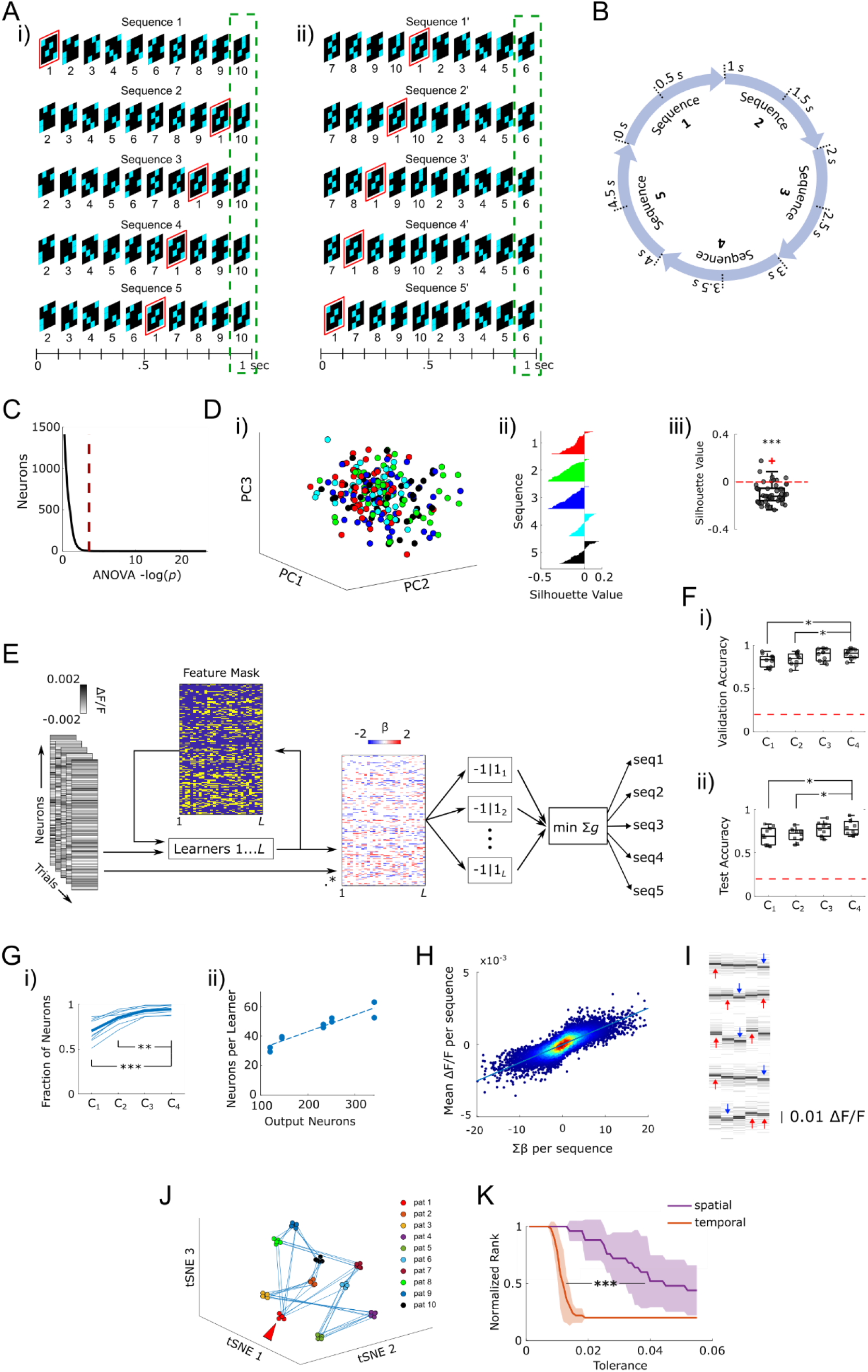
Temporal sequence inference. (**A**) Design of stimulation sequences: (i), pattern order is shown relative to pattern 10, during which output activity was examined (indicated by green dashed box), (ii) pattern order relative to pattern 6. Position of pattern 1 (indicated by red box) was changed between sequences. (**B**) Sequence delivery during 1 trial. (**C**) Histogram of *p* values, indicating significance of differences between sequences during stimulation with patterns 6 and 10, one-way ANOVA (*n* = 2190 stimulation responses, significance threshold indicated by red dashed line is *p* = 2.3*10^−4^). (**D**) (i) Representative graph of PCA scores of responses to pattern 6 in one culture, color-coded by sequence (*n* = 50 trials/sequence), and (ii) silhouette values for the same PCA analysis. (iii) Silhouette values averaged by sequence for patterns 6 and 10 in 5 cultures (*n* = 50). *** *p* < 0.001 on two-sided Wilcoxon signed rank test. (**E**) Classification pipeline. Output neuron activity population vectors were first used to identify useful features for each binary learner by RFE, resulting in a Feature Mask. Classifications of all *L* binary SVM learners (using *β* coefficients) for each observation (population vector) were combined via ECOC, with observation assigned to a sequence by minimizing aggregate loss (Σ*g*). (**F**) Multiclass classification accuracy: (i) validation accuracy of different ECOCs (C_1_, C_2_, C_3_, and C_4_), * *p* = 0.023 C_1_ vs. C_4_, and p = 0.046 C_2_ vs. C_4_, two-sample *t*-test, (ii) test accuracy, \**p* = 0.045 C_1_ vs. C_4_, and *p* = 0.05 C_2_ vs. C_4_, two-sample *t*-test. Validation and test accuracy of all ECOCs was significantly different from random level of 20% (indicated by red dashed line) with *p* < 0.001, two-sample *t*-test. (G) Utilization of output neurons: (i) Fraction of the total number of output neurons used by each ECOC. ** *p* = 0.002, *** *p* < 0.001, two-sample t-test. (ii) Number of neurons used per binary learner is linearly correlated to the number of output neurons per culture, Pearson correlation coefficient *r*^2^ = 0.89. *n* = 10 (2 patterns/culture, 5 cultures). (**H**) Neuron’s participation in sequence’s classification *(Σβ*) versus mean activity during that sequence, plotted as a density scatter plot (red color indicates higher density of points). Participation and activity are linearly correlated, Pearson’s *r*^2^ = 0.64, *p* < 0.001, *n* = 9959 participation-activity pairs. (**I**) Activity of 5 neurons with highest Σ*β* scores during evaluation pattern, for all 5 sequences. Light gray lines represent activity during sequence repeats, while thick black and dark gray lines represent mean +/-standard error of mean. Arrows represent positive (red) or negative (blue) contribution of activity to classification of the corresponding sequence. (**J**) Representative t-distributed stochastic neighbor embedding (t-SNE) of output neuron data averaged by sequence. Trajectories of output neuron activity during different sequences are indicated by blue lines. Red arrowhead indicates convergence of trajectories on output neuron state defined by spatial pattern 1 (pat 1), despite changing temporal position of pattern 1 during different sequences. (**K**) Spatial and temporal kernel quality represented by normalized response matrix rank calculated for different tolerances. Temporal rank drops at lower tolerances than spatial rank, *** p < 0.001, two-sample t-test, spatial *n* = 5 cultures, and temporal *n* = 10 (2 patterns/culture, 5 cultures).

We then determined whether temporal sequences could be inferred from culture activity despite lack of significance or separability using conventional statistics or dimensionality reduction. We employed recursive feature elimination (RFE) to minimize the number of features used to train the classifier. We designed custom Error-Correcting Output Codes (ECOCs) that combined different numbers of binary classifiers (learners) for multi-class classification into 5 sequences (Table S1). These ECOCs were termed C_1_, C_2_, C_3_, and C_4_. We reasoned that different sets of output neurons may have evoked activity that allows separation of a given pair of sequences; therefore, we carried out RFE for each binary learner. Binary learners were SVMs that produced *β* coefficients for classification. Output of all learners was combined using a hinge loss function to classify each output neuron population activity vector into 1 of 5 sequences (Fig. 4E). We partitioned pattern 6 and pattern 10 data for each culture into multiple training (90%) and testing (10%) sets. Binary learners used only data from output neurons identified by RFE (Fig. S6A). Number of output neurons used per learner was linearly correlated to number of output neurons in a culture (Fig. 4G(ii)), and ranged from 17% to 28% of total. Learners were trained using training data, and their individual training(validation) accuracy was determined using 8-fold cross-validation (Fig. S6B). Their test accuracy was then determined using the testing data set. Validation and test accuracies of multiclass classification into 5 sequences was determined using 20 random initiations of training and testing sets, respectively. Average accuracies for different ECOCs are shown in Fig. 4F (i ii). Median validation accuracy for best ECOC (C_4_) was 91%, and median test accuracy was 77%. Validation and test accuracies of all ECOCs were significantly higher than 20% random level. Best ECOC was characterized by its use of the highest proportion of neurons in cultures (∼95% of all output neurons, Fig. 4G(i), neuron utilization by ECOCs is shown in Fig. S6C).

We then sought to determine which aspects of output neuron activity were responsible for high classification accuracy. Linear SVM learners utilize *β* coefficients for binary classification. Number of *β* coefficients is equal to the number of features (post-RFE output neurons) utilized by a learner. Coefficients can be positive or negative, and their magnitudes determine relative contribution of a neuron to classification into ‘-1’ or ‘1’ class. We summed up *β* coefficients by neuron and by sequence (Σ*β*) for all learners for C_4_ ECOC, and found that each neuron could participate in classification of more than one sequence, with a positive contribution to some sequences, and negative contribution to others (Fig. S7). We then plotted the neuronal participation metric, Σ*β*, against the difference of that neuron’s mean activity during the corresponding sequence and mean activity during all sequences 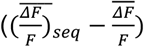 (Fig. 4H). We found that neuronal participation in classification of a sequence was significantly correlated to its mean activity during that sequence. To better understand this result, we plotted mean activity during all 5 sequences for 5 neurons with largest magnitudes of Σβ for each sequence in one of the cultures, and indicated each neuron’s participation in sequence classification (Fig. 4I). Relatively small differences in mean activity between sequences were sufficient to achieve high accuracy in classifying these sequences, despite relatively large variability between sequence repeats. We verified that sequence specificity was not a result of the dynamics of the calcium or jRGECO1a decay by calculating autocorrelation coefficient r of each culture’s output neurons. Autocorrelation beyond 100 msec was close to zero, indicating lack of memory in the fluorescence signal beyond that time (Fig. S8). We then compared encoding of spatial and temporal information. Trajectory of output neuron activity depended only on spatial input, returning to a state determined by spatial patterns regardless of temporal sequence (Fig. 4J). Kernel quality of the reservoirs, which determines their capacity for separating different inputs, can be measured by calculating reservoir matrix rank for different input classes (21). We found that the kernel quality of neuronal reservoirs for separating spatial information was significantly better than for separating temporal information; full rank was nevertheless achieved for separation of both types of information (Fig. 4K).

## Discussion

Our results show that activity of output neurons is substantially more influenced by spatial features of the input relative to temporal features (Fig. 3A vs. Fig. 4I). This is somewhat different than the concept of time processing proposed previously, where processing of spatial and temporal information was thought to be linked and encoded in a trajectory of a neural population through state space (7). State-space representation of our spatiotemporal sequence experiment is shown in Fig. 4J. State of the output neuron population is mostly determined by spatial features of the input. Even when a spatial pattern is delivered at different times in the sequence of patterns, output neuron population returns to nearly the same state, rather than evolving (pat1 in Fig. 4J). Instead of trajectory evolution, temporal features of the input were encoded in a subtle perturbation of the output neuron population activity, which is otherwise determined by spatial features of the input. Difference in spatial and temporal kernel quality of the reservoirs further supports this conclusion: reservoirs were able to separate different classes of spatial inputs at higher tolerances than temporal inputs, indicating that differences in output activity levels for different spatial inputs are higher than for different temporal inputs. Nevertheless, small perturbations were sufficient for reliable inference of temporal information.

Relatively small differences in activation of output neurons due to different sequences may be caused by a variety of mechanisms. These changes in somatic [Ca^2+^]_i_ may reflect small changes in the firing rate of output neurons. They may also reflect calcium transients due to synaptic activity. It is interesting to ask what is the mechanism by which the recurrently connected reservoir, formed by cultured dissociated neurons, encodes temporal information in these small [Ca^2+^]_i_ differences. Potential mechanisms include reverberant neuronal activity that stores temporal information in the evolving activation state of the network, or hidden processes such as short-term synaptic plasticity or long-lasting currents that store temporal information in the intracellular state of neurons. A limitation of this work is that we cannot distinguish between these mechanisms. While the state of neuronal activation was mostly determined by spatial input pattern, we could detect small differences in the state that were due to temporal input pattern features. These differences could represent small changes in the rate of reverberant activity. Alternatively, cellular processes such as short-term synaptic plasticity and spike frequency adaptation occur on a similar time scale as differences between sequences in this work (hundreds of milliseconds – seconds) (22–24), and may be responsible for temporal information encoding. It is also possible that both network and cellular mechanisms operate concurrently. One mechanism that we can rule out is intracellular calcium dynamics – our results show that decay time constant of [Ca^2+^]_i_ increase activated by stimulation with light patterns is ∼ 100-150 msec for output neurons, too fast to account for nearly a second-long memory exhibited by the reservoir.

A second limitation of this work is that we do not know whether small changes in output neuron activation that encode temporal information could translate to changes in activation of postsynaptic neurons in the (hypothetical) next processing layer. This question can be addressed in future work by construction of multi-layer, connected reservoirs that may be capable of hierarchical processing of various input features (25, 26). We can get an idea of the size and connectivity of the next processing layer from the relatively sophisticated readout function *f* (Fig. 1A(ii) and Fig. 4E) that was required to infer temporal information with high accuracy. Activity of each output neuron was evaluated by up to 30 different learners, with 2 separate pipelines for different ranges of delays. Biological neuron representation of such a readout function can thus be expected to consist of 50 – 100 neurons (assuming that a neuron can perform a function roughly equivalent to each learner), with each readout neuron connected to about a third of the output neurons in the reservoir. Connectivity between output reservoir and readout layer would thus be characterized by a relatively large fan-out (each output neuron has many postsynaptic partners) and large fan-in (each read-out neuron has many presynaptic partners). This kind of connectivity is characteristic of the intact cortex, suggesting that cortex possesses the architecture that enables hierarchical processing of temporally structured information by many separate readout functions.

Prior work on living neuronal reservoirs examined reservoir response to spatial (12) or mixed spatiotemporal inputs (13–15). In this work, we separated spatial and temporal information contained in the inputs, and showed that living neuronal reservoirs can convert purely temporal features of the input into a spatial code. In computational terms, collective activity of neurons in the living reservoir represented a spatial embedding of the temporal information in the inputs (27). This adds experimental evidence to prior body of theoretical work suggesting that reservoir computation may be a plausible mechanism for temporal->spatial conversion that occurs in sensory cortices (6, 7). High accuracy of inference of temporal information from spatial code suggests that the reservoir is capable of nearly complete retention of temporal information, at least at the time scale examined in this work. This property is enabled by rich neuronal intracellular dynamics or by reverberant activity, and may be harnessed for future development of computational devices that either contain living neurons (biohybrid devices) or artificial elements that mimic complex neuronal dynamics (neuromorphic devices).

## Conclusions

In summary, we found that reservoirs composed of living cortical neurons could convert temporal features of the input into reservoir state. State, represented by spatial pattern of activation of output neurons in the reservoir, contained information about inputs for at least 900 msec. Timing of input pattern occurrence could be determined with 100 msec precision. Accurate classification required the use of most output neurons, suggesting that timing information was encoded via population code. Possible mechanisms responsible for conversion of temporal information into spatial code include neuronal and synaptic dynamics, as well as reverberant activity of recurrently connected neurons in the reservoir. Reservoir computation may be a plausible mechanism for temporal/spatial code conversion that occurs in sensory cortices. High conversion fidelity, and relatively long memory of living neuron reservoirs may be harnessed in future biohybrid or neuromorphic computational devices.

## Materials and Methods

### Cell Culture

Polydimethylsiloxane (PDMS, Sylgard 184) films of 100 μm thickness were prepared by spin-coating 150 mm petri dishes. Wells were cut in the film using syringe needles. PDMS film with a well was then transferred to a cell culture-treated glass bottom dish which was coated with poly-D-lysine (PDL). Dissociated cortical neurons from post-natal day 0–1 Sprague-Dawley rat pups, Charles River Laboratories were prepared as described earlier (18). Approximately 2 μl of dissociated cell solution containing approximately 30 thousand cells were plated inside each PDMS well. For the first hour after plating cells, cultures were incubated at 37°C and 5% CO_2_ in a medium of 10% Fetal Bovine Serum (FBS) in Neurobasal-A supplemented with GlutaMAX and gentamicin (Fisher Scientific). After 1 hour, the medium was replaced with fresh culture medium (97.45% Neurobasal-A, 2% B27, 0.25% GlutaMAX, and 0.3% gentamicin) and the cultures were placed again inside the incubator. Half of the medium was replaced with fresh medium twice a week. All animal use protocols were approved by the Institution Animal Care and Use Committee (IACUC) at Lehigh University and were conducted in accordance with the United States Public Health Service Policy on Humane Care and Use of Laboratory Animals.

### ChR2 and jRGECO1a Expression

After 24 h post-plating, we replaced half of the culture medium with fresh culture medium containing 1 μl of Adeno-associated Viral 9 (AAV9) particles wih pAAV-hSyn-hChR2(H134R)-EYFP (at titer ≥ 1×1013 vg/mL) and 1 μl of AAV1 particles with pAAV.Syn.NES-jRGECO1a.WPRE.SV40 (at titer ≥ 1 × 1010 vg/ml) (both from Addgene). Neurons had significant ChR2 and jRGECO1a expression by day in vitro (DIV) 8 (Fig. 1B).

### Optical Detection of Neural Activity

Optical recordings were performed by placing the culture dish inside a mini-incubator on a stage of a dual-deck fluorescent inverted microscope (IX73, Olympus) starting from DIV 10 until DIV 20. Composition of artificial cerebrospinal fluid (ACSF) used as recording medium was, in mM: 140 NaCl, 2.4 KCl, 10 HEPES, 10 glucose, 2 CaCl_2_, 4 MgCl_2_, and 1 Na_2_HPO_4_. Atmosphere inside the mini-incubator was maintained at 37°C and 5% CO_2_. Changes in jRGECO1a fluorescence (580/610 nm excitation/emission) were captured at 20 frames per second (fps) except burst probability experiments when recordings were captured at 5 fps. To capture the change in intensity over time for individual neurons, Regions of Interest (ROIs) were manually defined for them in Fiji (ImageJ) software. Mean gray values of ROIs during each frame were measured and stored for analysis. Data analysis was performed in MATLAB (Mathworks). Fluorescence traces were converted to the ratio of change in fluorescence over background fluorescence (Δ*F/F*), with background fluorescence estimated via asymmetric least squares smoothing (28). Fluorescence ratios used for spatial and temporal inference analysis were filtered with a high pass 6^th^ order Butterworth filter with a cut-off frequency of 0.05 Hz to eliminate slow variability at time scales of 20 sec or greater.

### Patterned Optical Stimulation

Optical patterns were delivered by a pattern illuminator (Polygon 400G, Mightex), mounted on the second deck of the inverted microscope. Polygon 400G combines an externally-triggered light emitting diode (LED) source of 480 nm light with a digital micromirror device that enables spatial targeting of the light. We first optimized stimulation and recording parameters by recoding optically stimulated neurons in whole cell configuration (see Results and Supplementary Methods). We then set the illumination power at 10 mW/mm^2^ for all subsequent experiments. We divided the working area of Polygon 400G into 10×10, 20×20, or 30×30 grids with different spacing to create stimulation patterns for spatiotemporal inference experiments. Then we selected some of the grid cells for optical stimulation, and imported them into MATLAB to generate sequences of 10 patterns. Those sequences were then loaded into the Polygon 400G. Each pattern was illuminated by 5 pulses of light at 50 Hz with individual pulse-width of 10 msec. Each pattern was 100 msec long and each sequence consisting of 10 patterns was 1 sec long. Simultaneously with optical stimulation, we recorded changes in jRGECO1a fluorescence. Both Polygon 400G and the camera were synchronously triggered by an external source (Axon Digidata 1550B).

### Analysis of Evoked Activity

*Decay time constant τ*_*D*_. Neurons with significant activity during a pattern sequence had *p* < 0.05/neuron number on a two-sample *t*-test of maximum and minimum Δ*F*/*F*. Time for activity to decay from maximum Δ*F*/*F* to e^-1^ of maximum was defined as *τ*_*D*_ for that neuron.

*Spatial Spread*. Spatial spread of activity in output neurons was calculated per stimulation pattern. Positions of neurons with significant activity (*p* < 0.05/neuron number, two-sample *t*-test of activity during current and preceding patterns). Spatial spread was defined as the standard distance of active neurons, calculated by taking square root of sum of variance of neuron positions along *x* and *y* axes. Average spatial spread for all patterns in a culture was normalized by size of the field of view.

*Population activity vectors*. Population activity vectors for output neurons were used for spatial and temporal inference analyses. These vectors were constructed from individual neurons’ ΔF/F values during last 50 msec of a given pattern. A vector was constructed for each stimulation pattern repeat.

*Silhouette Value*. Silhouette value *s*_i_ for the *i*th point in PCA (representing PC scores of the *i*th population activity vector) were calculated as

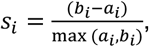

where *a*_i_ is the average distance from the *i*th point to the other points in the same cluster as *i* (i.e other responses to the same stimulation pattern), and *b*_i_ is the minimum average distance from the *i*th point to points in a different cluster (i.e. responses to different stimulation patterns), {*s*_*i*_ ϵ ℝ |−1 < *s*_*i*_ < 1}.

### Pattern Inference

Matrix of population activity vectors of selected output neurons was used as input for the classification learner toolbox of MATLAB. Matrix was constructed using vectors for all 10 stimulated patterns for one culture. Each vector was considered as a single observation, and assigned an appropriate class label (number of stimulation pattern associated with that vector). Observations were divided into a training set and a test set. Linear support vector machine (SVM) model was then trained using training data set, and validation accuracy (on classifying the training set) was determined using 5-fold cross validation. Test accuracy was then determined by classifying the data in the test set using the trained SVM model. Separate SVM models were generated for each culture.

### Temporal Sequence Inference

*Error-Correcting Output Codes (ECOC)* Inference of a temporal sequence required multi-class classification, with number of classes *K* = 5 (sequences). SVM models which we have utilized for spatial and temporal classification are binary, classifying observations into class 1 or -1. In order to apply them to a multi-class problem, ECOCs were used to reduce multiclass problem to a series of binary problems. A commonly used ECOC is one-versus-one coding design, processing output of *K*(*K*-1)/2 binary learners via a hinge error minimization function. This ECOC is the default setting of MATLAB multiclass classification algorithms, and was used for pattern inference. We found that its accuracy for classifying temporal sequences was not as good (Fig. 4G(i,ii), C_1_ is the one-versus-one ECOC). We therefore designed custom ECOCs C_2_ (10 learners) and C_3_ (20 learners), where each binary learner classified the observation into a set of 2 or 3 sequences, as well as ECOC C_4_ (30 learners) which combined C_1_ and C_3_ (Table S1). ECOC coding assigns an observation to a sequence that produces minimum collective loss of *L* binary learners among sequences *k*,

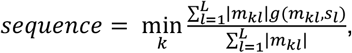

where *m*_kl_ is an element of ECOC matrix (*m*_kl_ ϵ {-1,1,0}), *s*_l_ is the classification score for learner *l* (s_l_ ϵ {-1,1}), and *g* is the loss function. We used hinge binary loss function of the form:

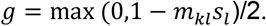

*Recursive Feature Elimination* (*RFE*). RFE was performed for each of the 30 possible binary learners defined by the largest custom ECOC (Table S1). Linear SVM classifiers were used as binary learners for both RFE and later ECOC-based multiclass classification. MATLAB ‘sequentialfs’ function was used to perform backward feature selection, or RFE. The function initially included all output neurons as features, and then sequentially excluded one feature with each iteration. Error of binary classification was determined for each iteration, and next feature to exclude was selected by the algorithm to minimize error. Iterations were continued until only one feature was left (Fig. S6A). Remaining features during an iteration with minimal error were used during subsequent learner training and multiclass classification (Fig. S6C).

### Software

Initial image and video analysis was performed using Fiji (ImageJ, NIH) to extract numerical data. Data analysis, including dimensionality reduction, inference, and statistics, was performed in Matlab R2023a (Mathworks).

### Statistics

Data generated by experiments was tested for normality using one-sample Kolmogorov-Smirnov test. Two-sample and multi-sample tests were chosen based on results of normality testing. Statistical tests and sample numbers are indicated in figure legends and text. Corrections of significance *p* values for multiple comparisons are also indicated as appropriate.

### Data Sharing

Matlab code and data used in this manuscript are available at: https://figshare.com/s/da3a975dd75838b034cd

## Acknowledgments

This research was supported by National Science Foundation (NSF) NCS grant ECCS 1835278 and Air Force Office of Scientific Research (AFOSR) grant FA9550-19-1-0419.

