## Supporting Information for "Temporal Information Encoding in Isolated Cortical Networks"

### **Supporting Information for**
### Temporal Information Encoding in Isolated Cortical Networks.

Zubayer Ibne Ferdous[1], Yevgeny Berdichevsky[1, 2, *].

1. Department of Electrical and Computer Engineering, Lehigh University, PA, USA.
2. Department of Bioengineering, Lehigh University, PA, USA.

*Corresponding author: Yevgeny Berdichevsky

**This PDF file includes:**

Supporting text
Figures S1 to S8
Table S1
SI References

**Other supporting materials for this manuscript include analysis code and data used to generate the figures, available at:** https://figshare.com/s/da3a975dd75838b034cd

#### Supporting Information Text

##### Supporting Methods

**Whole Cell Recording.** We replaced the culture medium with an artificial cerebrospinal fluid (ACSF) solution for performing electrophysiology. The ACSF solution contained (in mM): 140 NaCl, 2.4 KCl, 10 HEPES, 10 glucose, 2 $CaCl_2$, 1 $MgCl_2$, and 1 $Na_2HPO_4$ (pH 7.4)(1). Recordings were performed by placing cultures grown on coverslips on a perfusing chamber of an inverted microscope (Olympus). The chamber was kept at 37ºC using a temperature controller before placing the cultures. Electrodes for patch clamp were pulled from borosilicate pipettes (1.5 OD) using a pipette puller (Sutter instruments). The recipe of the pipette puller was tuned in a way so that the electrodes have 5-10 MΩ resistance when filled with an internal solution containing (in mM): 130 K-gluconate, 10 HEPES, 10 phosphocreatine, 5 KCl, 1 $MgCl_2$, 4 ATP-Mg and 0.3 mM GTP(2). All whole cell recordings were performed using Multiclamp 700B (Molecular Devices) amplifier. All whole cell recording signals were acquired at 10 kHz using Axon Digidata 1550B (Axon Instruments) and were saved as .abf files. Further analysis was done in MATLAB. All whole-cell recordings were performed starting from DIV 10 until DIV 20.

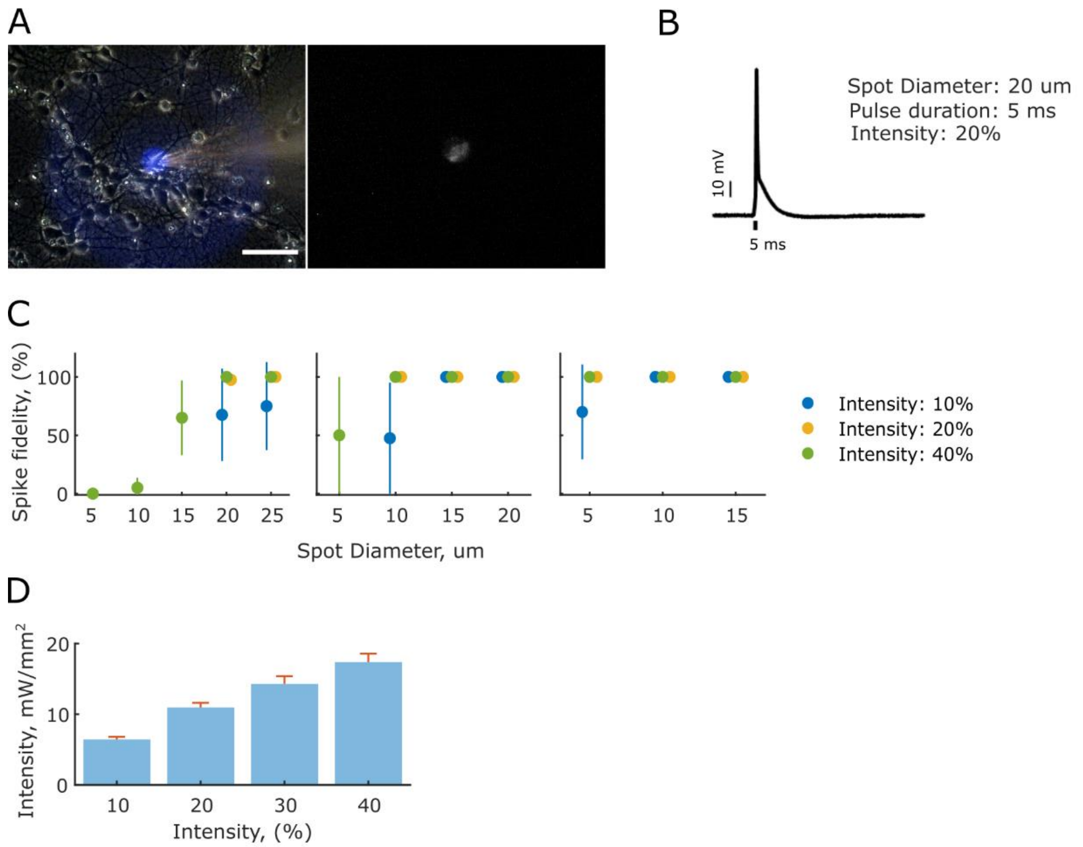


**Fig. S1.** Optimization of patterned stimulation parameters. (**A**) Combination of single neuron optical stimulation and whole cell recording. Phase (left) and fluorescence (right) micrographs show delivery of optical stimulus to the soma of a single, recorded neuron in the field of view. (**B**) Minimum tuning parameters such as the light spot size, pulse duration, and intensity for reliable evocation of action potentials in the recorded neuron. (**C**) Parameter tuning for reliable evocation of action potentials. Left, middle and right panels show action potential (spike) fidelity for light pulse durations of 2 msec, 5 msec and 10 msec, respectively. Means and standard deviation of 4 neurons from different cultures are shown. Each neuron received 10 optical stimuli. Minimum parameters shown in (B) were derived from this data. (**D**) Measurements of light power using a power meter at different intensity settings of the pattern stimulator. Means and standard deviations from 3 different spot sizes are shown.

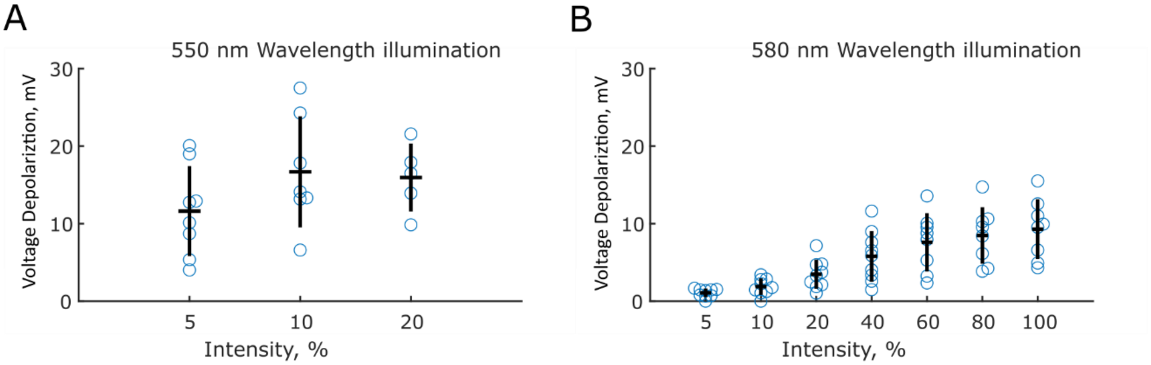


**Fig. S2.** Optimization of jRGECO1a excitation light wavelength and intensity. Voltage depolarization was measured when recorded neuron was illuminated with wide-field (**A**) 550 nm or (**B**) 580 nm wavelength light. Data was acquired from whole cell recordings from 8 neurons (550 nm light) and 9 neurons (580 nm light) from 8 different cultures on different DIVs. Black lines indicate mean and standard deviation.

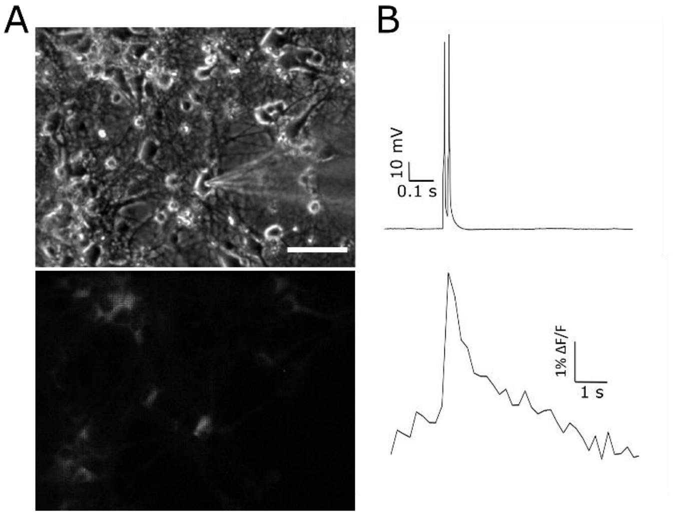


**Fig. S3.** jRGECO1a sensitivity with modified ex--citation light settings. (**A**) A neuron expressing jRGECO1a was stimulated by delivering current pulses through a whole cell recording pipette. (**B**) Representative simultaneous electrical (top) and optical (bottom) recordings are shown.

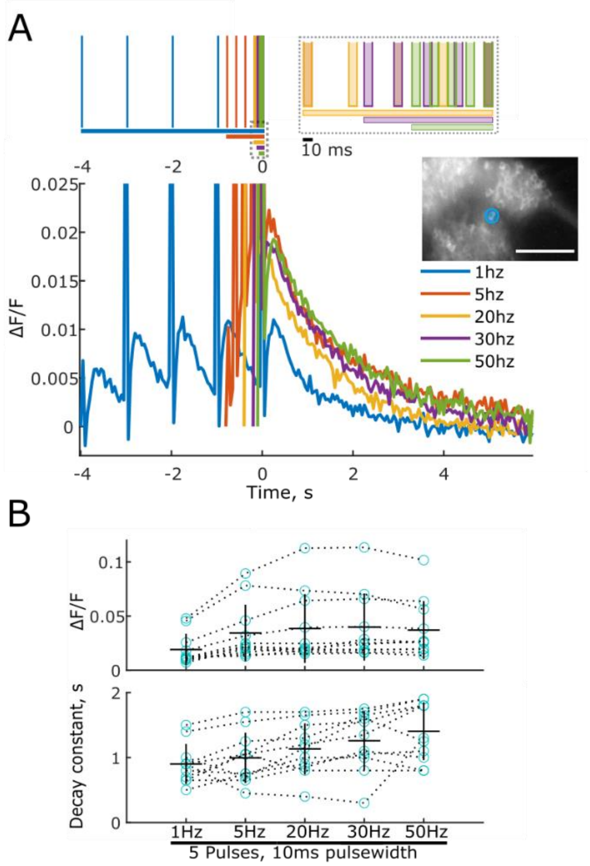


**Fig. S4.** Pulse train parameters. (**A**) Trains of 5 pulses of 10 msec duration were delivered to a single neuron (inset), with pulse frequencies ranging from 1 to 50 Hz. Top: durations of pulse trains of different frequencies. Bottom: representative response from a neuron to different pulse trains. (**B**) Maximum ΔF/F (top) and decay constant (from maximum ΔF/F) of neurons stimulated with pulse trains of different frequencies. Data for $n$ = 11 neurons from 11 cultures is shown; lines correspond to mean ± standard deviation. Maximum ΔF/F was not significantly different between 20 Hz and 50 Hz, and between 30 Hz and 50 Hz stimulation trains ($p$ = 0.91 and 0.83, respectively, two-sample $t$-test).

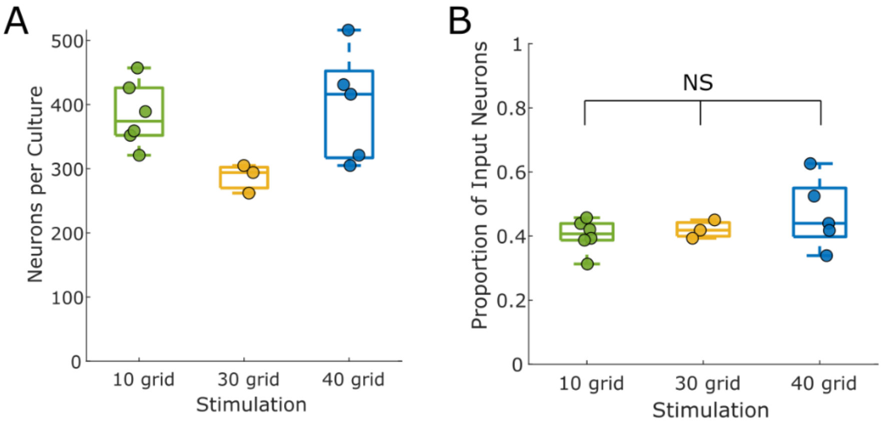


**Fig. S5.** Proportion of input neurons. (**A**) Number of neurons visible in the field of view for each culture used for grid size experiments. Boxes represent median, 25th, and 75th percentiles. Data for individual cultures is also shown. (**B**) Proportion of input neurons for stimulation patterns with different grid sizes. NS – no significant differences, one-way ANOVA $p$ = 0.36 (‘10 grid’ $n$ = 6, ‘30 grid’ $n$ = 3, ‘40 grid’ $n$ = 5 cultures).

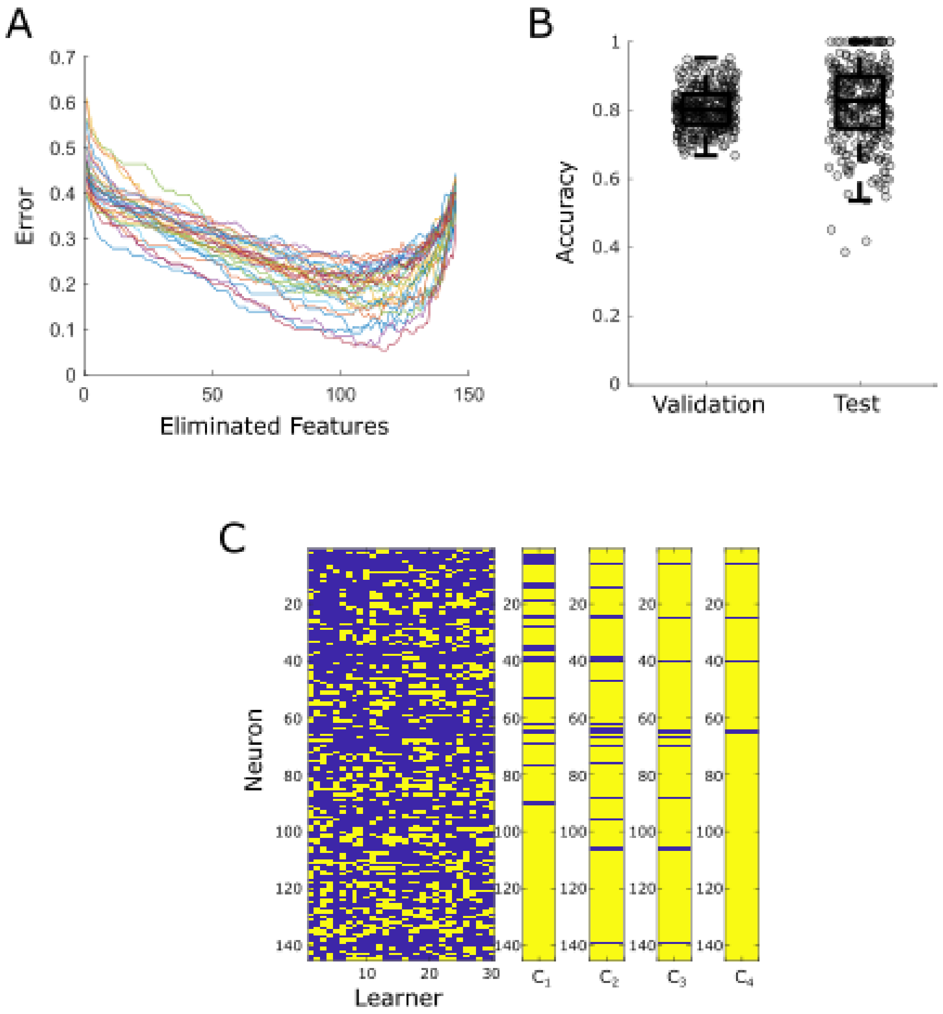


**Fig. S6.** Output neurons (features) used for sequence classification. (**A**) Representative plot of the change in binary classification error with recursive feature elimination for 30 learners (indicated by different colors) for one culture/evaluation pattern. (**B**) Validation and test accuracies for all binary learners used for 5 cultures and 2 evaluation patterns ($n$ = 300 learners), using optimum number of features identified by RFE. (**C**) Representative Feature Mask: neurons used by binary learners and by ECOCs for one culture/evaluation pattern. Used neurons are indicated by yellow color, unused neurons by blue color.

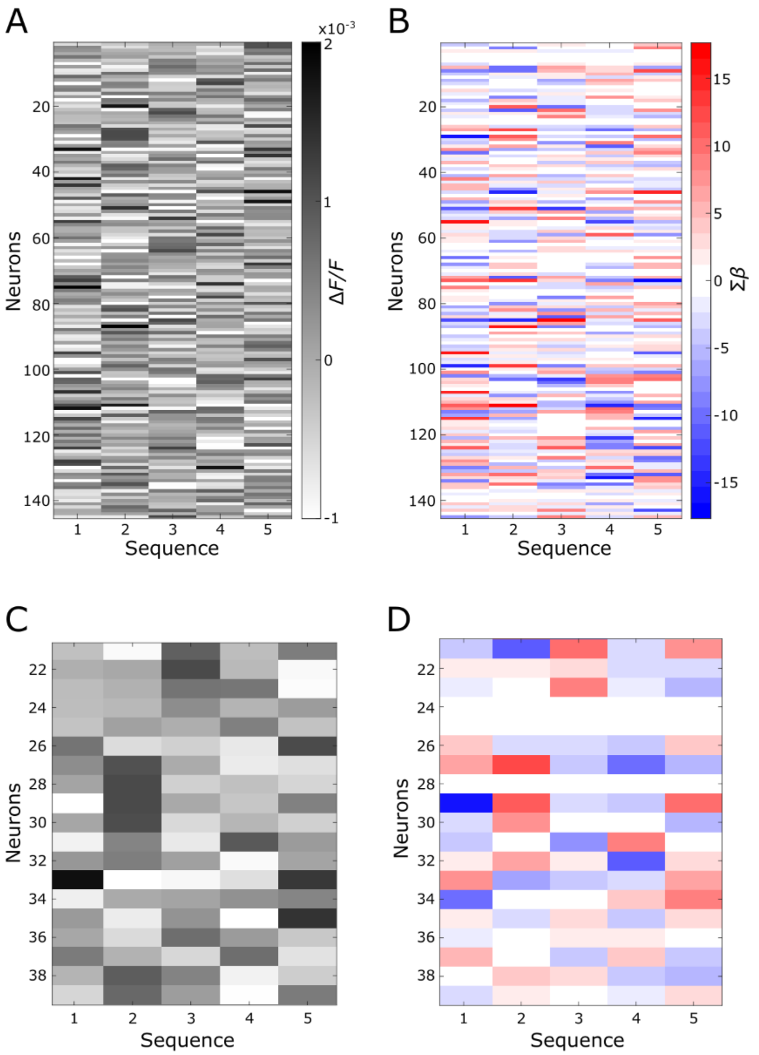


**Fig. S7**. Relationship between neural activity and participation in sequence classification, representative plots. (**A**) Average neural activity during 5 sequences, after subtracting all-sequence average, in one culture during one evaluation pattern. (**B**) Participation ($\Sigma\beta$) of neurons in (A) in sequence classification. (**C**, **D**) More detailed view of neurons 21-39 from plots in (A, B).

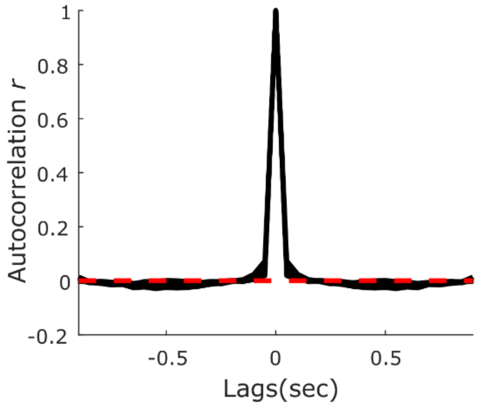


**Fig. S8.** Autocorrelation coefficient *r* of output neuron activity. Average normalized *r* for each culture is shown (*n* = 5 cultures, 5 sequences per culture). Red dashed line indicates *r* = 0.

| | $C_4$ | | | | | | | | | | | | | | | | | | | | | | | | | | | | | |
|---|---|---|---|---|---|---|---|---|---|---|---|---|---|---|---|---|---|---|---|---|---|---|---|---|---|---|---|---|---|---|
| | | | | | | | | | | | $C_3$ | | | | | | | | | | | | | | | | | | | |
| | $C_1$ | | | | | | | | | | | | | | | | | | | | $C_2$ | | | | | | | | | |
| **Learner #** | **1** | **2** | **3** | **4** | **5** | **6** | **7** | **8** | **9** | **10** | **11** | **12** | **13** | **14** | **15** | **16** | **17** | **18** | **19** | **20** | **21** | **22** | **23** | **24** | **25** | **26** | **27** | **28** | **29** | **30** |
| **Sequence 1** | 1 | 1 | 1 | 1 | 0 | 0 | 0 | 0 | 0 | 0 | 1 | 1 | 1 | 1 | 0 | 1 | 1 | 1 | 1 | 0 | 1 | 1 | 1 | 1 | 1 | 1 | 1 | 1 | 1 | 1 |
| **Sequence 2** | -1 | 0 | 0 | 0 | 1 | 1 | 1 | 0 | 0 | 0 | 1 | 1 | 1 | 0 | 1 | 0 | -1 | -1 | -1 | 1 | 1 | 1 | 1 | 1 | -1 | -1 | -1 | -1 | -1 | -1 |
| **Sequence 3** | 0 | -1 | 0 | 0 | -1 | 0 | 0 | 1 | 1 | 0 | -1 | -1 | 0 | 1 | 1 | -1 | 0 | 1 | 1 | -1 | 1 | -1 | -1 | -1 | 1 | 1 | 1 | -1 | -1 | -1 |
| **Sequence 4** | 0 | 0 | -1 | 0 | 0 | -1 | 0 | -1 | 0 | 1 | -1 | 0 | -1 | -1 | -1 | 1 | 1 | 0 | -1 | 1 | -1 | 1 | -1 | -1 | 1 | -1 | -1 | 1 | 1 | -1 |
| **Sequence 5** | 0 | 0 | 0 | -1 | 0 | 0 | -1 | 0 | -1 | -1 | 0 | -1 | -1 | -1 | -1 | -1 | -1 | -1 | 0 | -1 | -1 | -1 | 1 | -1 | -1 | 1 | -1 | 1 | -1 | 1 |

**Table S1.** Error Correcting Output Codes (ECOCs). Binary learners 1-30 used by ECOCs $C_1$ (10 learners), $C_2$ (10 learners), $C_3$ (20 learners), and $C_4$ (30 learners) to perform binary classification of a subset of sequences 1-5.
